# CellPatch: a Highly Efficient Foundation Model for Single-Cell Transcriptomics with Heuristic Patching

**DOI:** 10.1101/2024.11.15.623701

**Authors:** Hanwen Zhu, Yushun Yuan, Jiyuan Yang, Kangwen Cai, Nana Wei, Senxin Zhang, Lu Wang, Wen-Jie Jiang, YuanChen Sun, An Liu, Futing Lai, Yu-Juan Wang, Zeyu Ma, Xiaoqi Zheng, Hua-Jun Wu

## Abstract

The rapid advancement of foundation models has significantly enhanced the analysis of single-cell omics data, enabling researchers to gain deeper insights into the complex interactions between cells and genes across diverse tissues. However, existing foundation models often exhibit excessive complexity, hindering their practical utility for downstream tasks. Here, we present CellPatch, a lightweight foundation model that leverages the strengths of the cross-attention mechanism and patch tokenization to reduce model complexity while extracting efficient biological representations. Comprehensive evaluations conducted on single-cell RNA-sequencing datasets across multiple organs and tissue states demonstrate that CellPatch achieves state-of-the-art performance in downstream analytical tasks while maintaining ultra-low computational costs during both pretraining and finetuning phases. Moreover, the flexibility and scalability of CellPatch allow it to serve as a general framework that can be incorporated with other well established single-cell analysis software, thereby enhancing their performance through transfer learning on diverse downstream tasks.

## Main

Single-cell RNA sequencing (scRNA-seq) technology has revolutionized molecular biology in recent years through its capacity to generate high-resolution transcriptomic profiles at unprecedented cellular scales^1–3^. This technological advancement enables precise measurements of gene expression patterns in individual cells, yielding crucial insights into cellular heterogeneity, developmental trajectories, and disease mechanisms^4,5^. Meanwhile, The establishment of the Human Cell Atlas^6^ has further facilitated the systematic study of gene regulatory networks at single-cell resolution. Deep learning approaches have emerged as powerful tools for integrating and modeling large-scale scRNA-seq datasets, revealing complex cellular states and dynamics^7,8^. Recent Transformer-based foundation models of cells, such as scBERT^9^, scGPT^5^, GeneFormer^10^, scFoundation^11^, and LangCell^12^, particular show promise in effective feature extraction and versatile downstream applications, providing new insights into cellular functions. These models demonstrate that Transformer-based architectures^13^ significantly outperform traditional deep learning algorithms, including scVAE^14^ and scVI^15^, in learning representations of single cells. However, despite their success in processing single-cell data and addressing downstream analytical tasks^16^, current foundation models face significant challenges. The high dimensionality of transcriptomics data introduces considerable computational complexity within attention-based architectures, while batch-specific variations in gene detection necessitate complex preprocessing workflows. These challenges collectively constrain the models’ transferability, scalability, and flexibility across a range of biological applications.

To address these challenges, we present CellPatch, a novel foundation model that employs an effective gene patching strategy to reduce model complexity. Unlike text sequences, which can be chunked based on syntactic order, or images, where pixels can be patched according to spatial position, there is currently no universal patching strategy specifically designed for gene expression data^17^. CellPatch employs an innovative cross-attention mechanism where integrated patch tokens automatically patch genes as prior information and extract patch-level features. CellPatch is designed to directly execute downstream tasks using an encoder coupled with a task-specific decoder. Through comprehensive evaluation using extensive real datasets, CellPatch demonstrates superior performance compared to state-of-the-art (SOTA) algorithms. Moreover, the modular design of CellPatch allows for integration with existing SOTA models to enhance their capabilities. When integrated with STAGATE^18^, a prominent spatial transcriptomics analysis algorithm, CellPatch significantly improves spatial domain identification, leading to more precise hierarchical structural determinations.

The core innovation of CellPatch lies in its distinctive pretraining method and feature extraction mechanism (Fig. 1a). CellPatch extracts patch features from scRNA-seq data through unsupervised masked gene modeling (Methods), utilizing a comprehensive dataset of 10M cells (described in Data Availability). To optimize computational efficiency, we retained only non-zero elements, with each assigned a gene token that encodes positional information. These elements are subsequently integrated into input features through a sophisticated combination of Patch Tokens and cross-attention mechanisms (Fig. 1b). This approach enables CellPatch to extract gene expression features across diverse cellular contexts and generate standardized patch features of fixed dimensionality. The subsequent application of the self-attention mechanism to these patch features facilitates deep feature extraction, yielding higher-quality representations. A key innovation in the decoder architecture is the incorporation of gene embeddings as semantic prompts, which are not only utilized within the encoder but are also reintroduced in the decoder for enhanced contextual integration. By utilizing the cross-attention mechanism, the model effectively extracts information from patch features and accurately reconstructs gene expression profiles through the decoder. This dual-phase application of gene embeddings enhances the semantic representation capabilities of the gene embeddings.

**Fig. 1.**
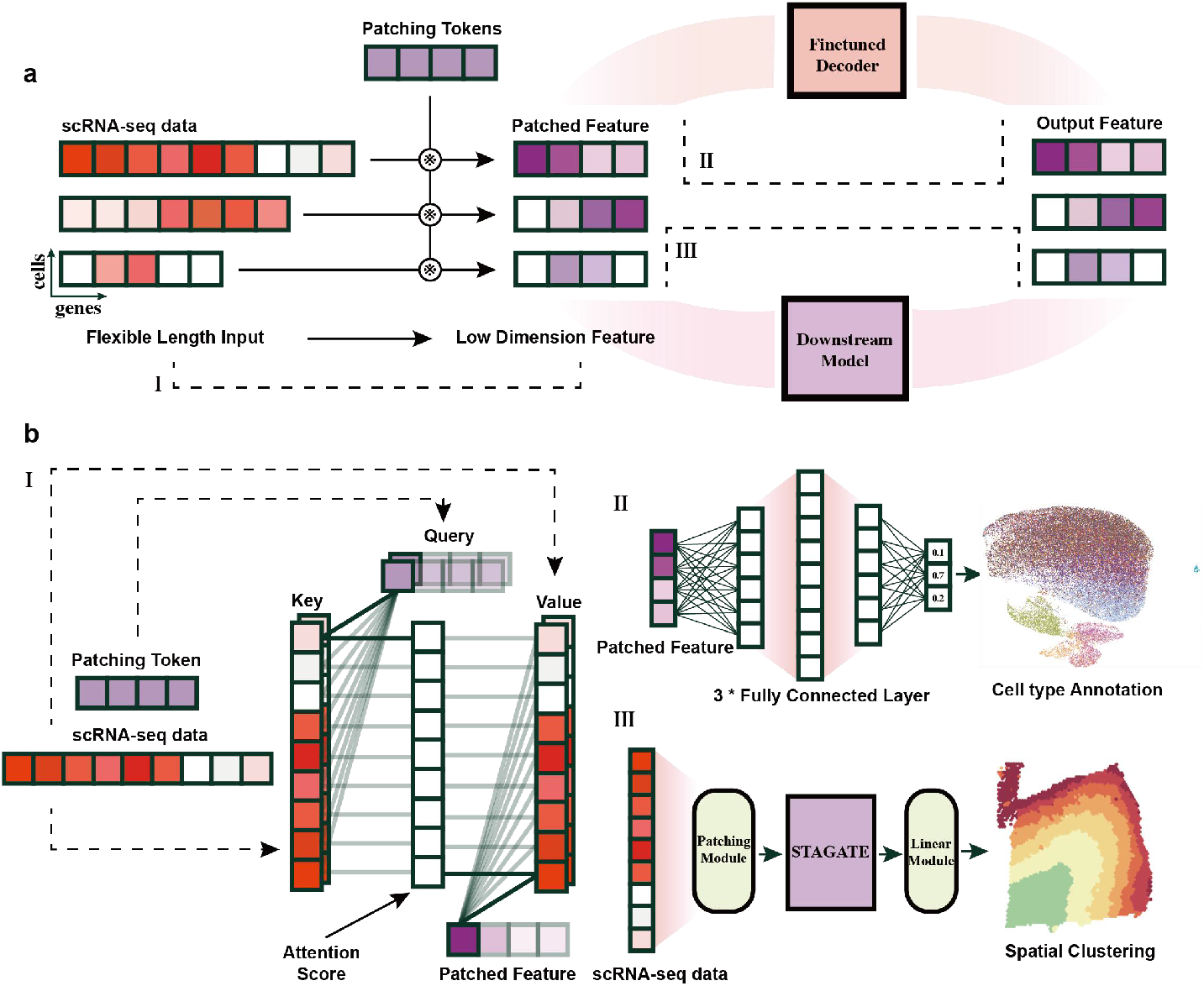
Model Framework. **a**, Overall framework of CellPatch. CellPatch reduces the dimensionality of single-cell RNA sequencing data by utilizing patch tokens obtained through pretraining, enabling application to various downstream tasks via fine-tuning or embedding. **b**, I Patch Module. Illustrating how a single cell derives patch features through the cross-attention mechanism. II Cell type annotation. Cell features are classified through three dense layers. III Model extension. The patching module and an additional fully connected layer are integrated into the existing STAGATE framework.

To showcase the embedding capability of CellPatch, we evaluated its cell type annotation performance against competitive algorithms across diverse scRNA-seq datasets that include a variety of organs and tissue states. Taking Zheng68K dataset as an example, we visualized the prediction results of CellPatch (Fig. 2a) and annotated cells that were not consistently predicted. We further calculated the confusion matrix to assess classification consistency (Supplementary Figure 1). Model performance was systematically assessed using accuracy and F1 score metrics. CellPatch demonstrated superior performance across all benchmark datasets compared to existing methods, achieving higher accuracy (Fig. 2b) and F1 scores (Supplementary Figure 2). Additionally, we calculated the runtime required by different models to perform the annotation task by incrementally increasing number of genes per cell as input (Supplementary Figure 3). CellPatch excelled in both speed and tolerance for higher number of input genes, outperforming other models and demonstrating its efficiency, stability, and broad applicability across various tissue states and experimental platforms.

**Fig. 2.**
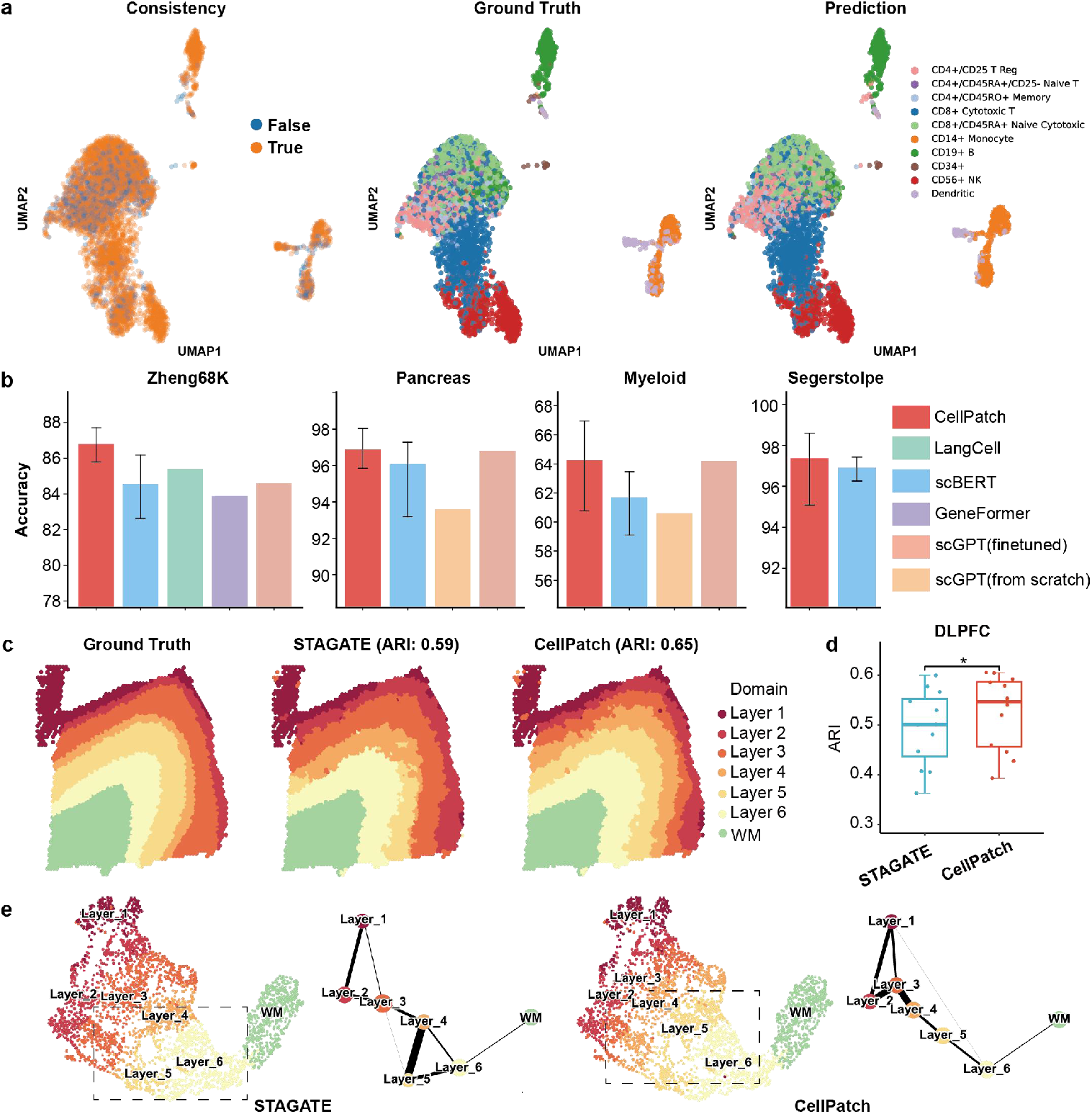
Performance of CellPatch on scRNA-seq and Spatial Transcriptomic Dataset. **a**, The cell type prediction performance of CellPatch on Zheng68K dataset. (Left) Consistency between the ground truth and the predicted results of CellPatch; (Middle) UMAP visualization of the ground truth; (Right) UMAP visualization of the predicted results. **b**, Cell type annotation accuracies of various methods on real scRNA-seq datasets, including Zheng68K, Pancreas, Myeloid, and Segerstolpe. Error bars indicate 95% confidence intervals across 10 replicates. **c**, Clustering results of original STAGATE and CellPatch+STAGATE on sample 151675 from the DLPFC dataset. The ground truth for tissue regions is based on the manual annotation of six cortex layers or white matter (WM) provided by the original study. **d**, Boxplots illustrating clustering accuracies across all 12 sections for original STAGATE and CellPatch+STAGATE. Asterisk (*) denotes p-value less than 0.05 based on a paired one-sided t-test. **e**, UMAP visualizations and PAGA graphs of sample 151675 generated from original STAGATE and CellPatch+STAGATE embeddings, respectively.

To demonstrate the transfer learning capacity of CellPatch, we used pretrained CellPatch model to assist STAGATE in capturing transcriptional heterogeneity with spatial context. We assessed the spatial domain detection performance of CellPatch combined with STAGATE (CellPatch+STAGATE) and the original STAGATE using the human dorsolateral prefrontal cortex (DLPFC) data by 10X Visium. In the following section, CellPatch+STAGATE is referred to as CellPatch, and the original STAGATE is referred to as STAGATE. In a representative analysis of slice 151675, which comprises 3,592 spots across 33,538 genes annotated into six neuronal layers and white matter, CellPatch effectively delineates distinct cortex layers and accurately recovers layer boundaries. In contrast, STAGATE struggles to accurately recover cortical layers 4 and 5, resulting in inferior clustering performance (STAGATE: ARI 0.59, CellPatch: ARI 0.65) (Fig. 2c). Moreover, CellPatch significantly enhances clustering performance across the twelve samples compared to STAGATE (Fig. 2d). Manual annotations and clustering results for other DLPFC slices are shown in Supplementary Figure 4. Furthermore, we performed UMAP and Partition-based graph abstraction (PAGA^19^) graph visualization of low-dimensional embeddings generated by CellPatch and STAGATE. CellPatch depicts the spatial trajectory from layer 1 to layer 6 and white matter in the UMAP plot, as well as in the PAGA graph, reflecting the established “inside-out” pattern of corticogenesis. However, since layers 4 and 5 cannot be effectively separated, the PAGA graph of STAGATE embedding reveals a circular spatial trajectory pattern from layer 4 to layer 6 (Fig. 2e). Gene set enrichment analysis (GSEA^20,21^) revealed layer-specific enrichment of key pathways, including myelin sheath, cytoplasmic translation, and synapse-related pathways, highlighting critical transcriptional programs in cortical development (Supplementary Figure 5). These findings were further validated using the STARmap dataset with single-cell resolution (Supplementary Figure 6). In summary, these results demonstrated CellPatch’s capacity to enhance the performance of downstream analysis methods through transfer learning.

To investigate the interpretability of CellPatch, we analyzed the correlations between attention scores from the first-layer patching process and gene embeddings. Using Zheng68k dataset, we extracted and normalized the attention weights during the transformation of gene expression data into patch features. The normalized attention score matrix revealed distinct cell type-specific patterns (Supplementary Figure 7). For example, in CD8+ Cytotoxic T cells, we observed elevated attention scores for the *IL32* gene, a gene known to trigger cytokine production, including *TNFα*, which aligns with the biological role of CD8+ Cytotoxic T cells in producing host defense cytokines. On the heatmap, we highlighted representative cell type-specific markers, demonstrating that the model’s attention scores were significantly higher for these genes within their respective cell populations, indicating a targeted recognition of distinct cellular phenotypes.

Furthermore, we investigated whether the model-learned gene embeddings contained semantic information beyond merely providing positional context for expression values in downstream tasks. We extracted the top 30 marker genes for each cell type in Zheng68k dataset, along with marker genes reported in the original publication, and generated UMAP visualizations of their embedding weights (Supplementary Figure 8). The analysis revealed remarkable patterns, with embedding weights logically representing gene-gene relationships. For example, multiple genes involved in cytotoxic pathways expressed in CD56+ NK cells clustered near *GZMK* in the lower left region, which was identified as an NK cell type-specific gene in original Zheng68k study. Similarly, platelet-associated genes *PF4* and *PPBP* clustered in the upper region, which also contained multiple CD34+ cell type-specific marker genes. These analyses demonstrate that CellPatch achieves cell type-specific recognition while effectively learning gene-gene interaction relationships, validating both its robust learning capability and post-training interpretability.

In summary, the comprehensive experimental results presented herein validate the effectiveness of CellPatch across diverse single-cell transcriptomic and related data types, establishing a robust foundation for its further utilization. The model demonstrates exceptional interpretability and exploration potential in both cell type classification and gene relationship analysis, highlighting the advantages of the CellPatch architecture. While our current analyses are based on training results from 10M data points over 50 epochs, UMAP visualization of the pre-training results suggests potential for further model optimization. By increasing training iterations, and broadening the sample space through additional data collection, CellPatch could be a more comprehensive single-cell foundation model. In conclusion, the CellPatch architecture provides an efficient training paradigm for gene-centric data analysis, addresses the gap in scRNA-seq data patching, and achieves excellent performance across multiple aspects of single-cell analysis.

## Methods

### CellPatch overview

CellPatch introduces the concept of patch tokens, which extract features from input gene expression data through a cross-attention mechanism. Through pretraining, we obtained a set of patch tokens with high generalization capability, effectively reducing the dimension of gene expression data to a unified meta-gene level. CellPatch improves upon masked gene modeling (MGM), a commonly used training strategy, by extending the utility of gene embeddings: beyond their conventional role as positional encodings in the encoder, they are also employed as prompts in the decoder for expression value reconstruction, thereby enriching their semantic representation.

### Gene and expression embedding

CellPatch employs a distinctive embedding approach tailored to the unique structure of single-cell data. Since each expression vector contains mixed information of both gene IDs and their corresponding expression values, mapping these components into feature vectors constitutes a critical step. In this study, we assigned a learnable feature vector to the *i*-th gene:

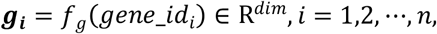

where ***g***_*i*_ represents the gene embedding vector, *f*_*g*_ denotes the gene-to-embedding transformation layer, *dim* denotes the embedding dimension, and *n* denotes the number of genes. This maps gene IDs into a predefined feature space. For gene expression values, we implemented a robust processing approach. We initially performed standard normalization on the data. Following the methodology employed in scBERT, the expression values are then clipped at a maximum of 5 and rounded to discrete values before being fed into the model. Subsequently, these processed values are transformed through an embedding layer into *dim*-dimensional expression features

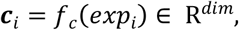

where ***c***_*i*_ represents the count embedding vector, *f*_*c*_ denotes the count-to-embedding transformation layer, and *exp*_*i*_ represents the count value of gene *i*.

Finally, the feature representation for a single cell at input is denoted as

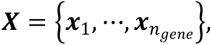

where ***x***_*i*_ = ***g***_*i*_ + ***c***_*i*_. This design not only effectively captures information about both gene ID and expression value, but also provides rich contextual information for the model, thereby enhancing performance in downstream tasks.

### Pretrain Encoder

In the encoder design of CellPatch, we proposed a novel architecture that differs from traditional Transformer structures. The model predefines a set of Patch Tokens and generates patch features by extracting information from cell features through cross-attention mechanism. Specifically, we first defined a set of ***PT*** (Patch Tokens) as

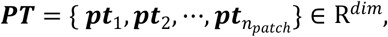

where *n*_*patch*_ represents the number of patch tokens, a hyperparameter that controls the dimension reduction ratio of gene features to patch features. The cross-attention blocks (Supplementary Note 1) then transform the input cell features ***X*** into patch features ***PF*** (patch feature), which is formulated as

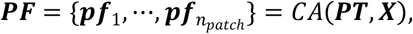

where *CA* denotes Cross-Attention blocks. Subsequently, these patch features are further processed through self-attention blocks (Supplementary Note 2) to obtain the final patch feature representation ***PF***^**′**^, which serves as input for various downstream tasks, formulated as

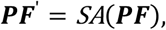

where *SA* denotes Self-Attention blocks.

### Pretrain Decoder

For the pretraining decoder design of CellPatch, we developed a prompt-based cell reconstruction architecture. We innovatively utilized gene embeddings as decoder prompts to reconstruct their corresponding gene expressions. This process extracts information from ***PF*** ^**′**^ through cross-attention mechanism and reconstructs gene expression via self-attention modules. Specifically, the reconstruction process can be formulated as

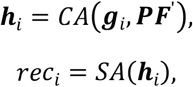

where ***h***_*i*_ represents the intermediate features extracted from ***PF*** ^**′**^ using gene embedding ***g***_*i*_ as the query in cross-attention operations, and *rec*_*i*_ denotes the reconstructed expression value for the *i*-th gene. The cross-attention block and self-attention blocks consist of multi-head attention layers followed by layer normalization and feed-forward networks.

During pretraining, we employed mean squared error (MSE) as the loss function to measure the discrepancy between the reconstructed expression values *rec*_*i*_ and the original expression values *c*_*i*_, which can be formulated as

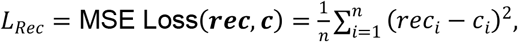

where *n* denotes the number of gene prompts input to the decoder for each cell. This reconstruction loss guides the model to learn meaningful representations of gene expression patterns through the pretraining process.

Our experimental results demonstrate that, as the number of training iterations increases, the intermediate representations ***PF*** ^**′**^ generated by the model progressively develop the ability to discriminate between different cell types, validating the effectiveness of our training approach (Supplementary Figure 9).

### Downstream task Classification Task Decoder

For cell type annotation tasks, we implemented a classification decoder architecture that comprising stacked dense layers to predict cell type assignments. Specifically, we first flattened the learned intermediate representations ***PF*** ^**′**^, which are then processed through a Dense Block consisting of three fully connected layers with dimensionality reduction.

The Dense Block projects the flattened features into a probability distribution over the predefined cell type space for each input cell. This transformation can be formulated as

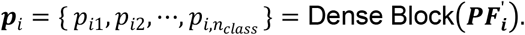

To quantify the performance, we adopt the cross-entropy criterion as our loss function, which measures the dissimilarity between the predicted probability distribution and ground truth annotations. The loss function is defined as

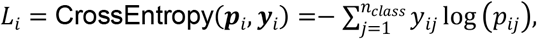

where 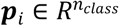 denotes the predicted probability distribution for the *i*-th cell, *n*_*class*_ represents the cardinality of the cell type set, and ***y***_*i*_ is the one-hot encoded ground truth label.

### Downstream Application: Enhancing Spatial Transcriptomics Analysis through CellPatch-STAGATE Integration

To further demonstrate the extensibility of CellPatch and its potential for complex downstream tasks, we proposed an innovative integration with STAGATE, a prominent spatial transcriptomics analysis framework.

Specifically, we utilized CellPatch to process spatial transcriptomic data and generate ***PF***, which is then concatenated with the original input features to serve as the enhanced input ***x***_*i*_ for the modified STAGATE module. The downstream analysis is performed through the intermediate representations ***z***_*i*_, which can be formulated as

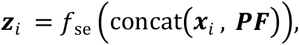

where *f*_*se*_ denotes the encoder of STAGATE. Given the graph autoencoder architecture of STAGATE, we added an additional Dense layer at the end of decoder to maintain consistency with the original feature space

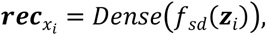

where 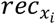 denotes the reconstructed features and *f*_*sd*_ denotes the decoder of STAGATE.

### Implementation details Gene Symbol Unification

We established a comprehensive gene reference set based on the CELLxGENE dataset, encompassing 60,690 genes along with their corresponding Ensemble IDs and gene symbols. This reference set incorporates, but not limited to human protein-coding genes and common mitochondrial genes. During the training and fine-tuning processes, each gene is assigned a unique token ID. For genes not present in the training set, new token IDs are dynamically allocated to ensure that no critical information is lost during model adaptation.

### Data Preprocessing

We implemented a standardized data preprocessing pipeline using the *Scanpy* framework, which includes quality control filtering, library size normalization (scaling total counts per cell to 1e4), and log transformation (log1p) of normalized counts. For the pretraining phase, we further optimize the input by filtering out zero-expression genes to reduce computational complexity, followed by padding the sequences to a uniform length to facilitate efficient batch processing.

### Differential gene expression analysis and spatial domain annotation

We employed the *MAST* algorithm implemented in the *FindMarkers* function of the *Seurat* package to identify differentially expressed genes for each spatial domain with a 5% FDR threshold (Benjamin-Hochberg adjustment). Then, Spatial domains are annotated by marker genes and comparing expression spatial patterns against manual annotations.

### Trajectory inference

After obtaining the clustering labels, we employed the PAGA algorithm to depict spatial trajectory.

### Gene enrichment analysis

We performed gene set enrichment analysis (GSEA) on the differentially expressed genes sorted by adjusted p-values using *enrichGO* function in the *clusterProfiler*^22^ package, showing the top five enriched pathways. Gene sets are downloaded from the Molecular Signatures Database (MSigDB, Broad Institute) including C2 (KEGG), C5 (GO BP: biological process, GO CC: cellular component, GO MF: molecular function).

### Evaluation metrics

To quantitatively assess the cell type annotation performance, we employed two complementary metrics: accuracy and F1 score. The accuracy metric, defined as the ratio of correctly classified cells to the total number of cells, is formulated as:

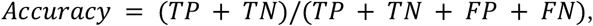

where *TP, TN, TP*, and *TN* denote true positives, true negatives, false positives, and false negatives, respectively.

To provide a more comprehensive evaluation, particularly for imbalanced cell type distributions, we additionally utilized the F1 score, which is the harmonic mean of precision and recall

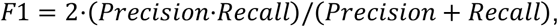

where

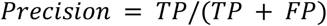

and

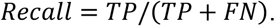

We adopted adjusted Rand index (ARI) to measure clustering performance of CellPatch and STAGATE in spatial domain detection. Specifically, given two sets of clustering labels, the ARI is calculated as

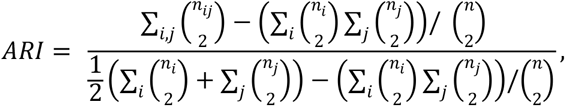

where *n*_*ij*_ is the number of spots overlapped by cluster *i* and cluster *j. n*_*i*_ and *n*_*j*_ are the number of spots in cluster *i* and *j*, respectively.

### Model Parameters Pretrain

The model is implemented using *PyTorch*. The model architecture is based on a Transformer structure with 64 patch tokens and an embedding dimension of 32. The encoder consists of 1 cross-attention layer followed by 2 self-attention layers for pathway embedding processing, with 2 attention heads. The encoder projects the output features into a 10-dimensional space. The decoder comprises 1 cross-attention layer and 1 self-attention layer for gene embedding processing, with final projection to 1-dimensional output. We set the maximum number of genes to 70,000 and utilized 64 learned pathway tokens as preset embeddings.

For training, we used the Adam optimizer with a learning rate of 1e-4 and weight decay of 5e-5. The model was trained for 50 epochs with a batch size of 512. Input gene expression data underwent log transformation, and random masking was applied to 30% of non-zero values during training. The maximum sequence lengths were set to 3,000 and 1,000 for encoding and decoding respectively.

### Cell type annotation

For the downstream cell type annotation task, we finetuned the pretrained model by replacing the original decoder with three dense layers. The decoder architecture consists of a first dense layer that projected the pathway embeddings (dimension: *n*_*pathway*_ × 32) to 512 dimensions, followed by a second layer reducing to 100 dimensions, and a final classification layer outputting predictions for cell types. Each of the first two layers is followed by ReLU activation and dropout regularization. The model is finetuned for 50 epochs using a batch size of 256. The Adam optimizer was employed with a learning rate of 1e-3 and weight decay of 5e-4. Gradient clipping was applied with a threshold of 5 to prevent exploding gradients.

### CellPatch-STAGATE Integration

The first layer of CellPatch is integrated into the STAGATE framework to enhance spatial feature extraction. The model architecture maintains the original cross-attention layers for patch-gene interactions, followed by STAGATE’s graph attention network with a two-layer structure to obtain 30-dimensional latent representations ***z***_*i*_. Finally, the pathway embeddings are projected through a linear decoder to match the output dimension.

The integrated model is trained for 1,000 epochs using the Adam optimizer with a learning rate of 1e-4 and weight decay of 1e-4. Gradient clipping was applied with a threshold of 0.1 to ensure stable training.

## Supporting information

Supplemental Figures and Notes for CellPatch

## Data availability

All data used in this study are publicly accessible. For pretraining, a total of 10M cells were sampled from the CELLxGENE database (https://cellxgene.cziscience.com/datasets), which comprises 30M cells across various cell types. Our model was evaluated using multiple publicly available datasets, including Zheng68k dataset from the ‘Fresh 68K PBMCs’ section (https://support.10xgenomics.com/single-cell-gene-expression/datasets)^23^, pancreatic datasets from the scGPT foundation model processed datasets (https://hemberg-lab.github.io/scRNA.seq.datasets/)^24^, the Myeloid dataset from Gene Expression Omnibus (GEO: GSE154763)^25^, and the Segerstolpe dataset from ArrayExpress (E-MTAB-5061)^26^. The human DLPFC dataset measured by 10X Visium is available at (http://research.libd.org/spatialLIBD/)^27^. The STARmap dataset for mouse medial prefrontal cortex dataset is available at (https://www.starmapresources.org/data)^28^; Data and scripts associated with this study are available at Github. (https://github.com/HanwenZhu98/CellPatch).

## Code availability

The source code of CellPatch is freely available on Github (https://github.com/HanwenZhu98/CellPatch).

## Acknowledgements

This work was supported by the National Natural Science Foundation of China (32270683 and 32470662), the Beijing Natural Science Foundation (5242006), the Fundamental Research Funds for the Central Universities (BMU2021YJ064 and PKU2022LCXQ027), and the National Key R&D Program of China (2021YFC1712805) to H.J.W.; the National Natural Science Foundation of China (61972257), the Natural Science Foundation of Shanghai (20JC1413800), and the National Key R&D Program of China (2018YFA0900600) to X.Z.. We gratefully acknowledge the High-performance Computing Platform of Peking University for conducting the data analyses.

## Author information

## Ethics declaration

No ethical approval was required for this study. All utilized public datasets were generated by other organizations that obtained ethical approval.

## Competing interests

The authors declare no competing interests.

## Supplementary information

Evaluation Metrics & Supplementary Figures

## Notes

### Competing Interest Statement

The authors have declared no competing interest.

