## Supplemental Figures and Notes for CellPatch for "CellPatch: a Highly Efficient Foundation Model for Single-Cell Transcriptomics with Heuristic Patching"

**Supplementary Notes**

**Supplementary Node 1. Cross-attention Mechanism**

Let $\boldsymbol{X}\boldsymbol{\in}\boldsymbol{R}^{\boldsymbol{m\times n}}$ denote the input feature vectors, where $\boldsymbol{m}$ represents the sequence length and $\boldsymbol{n}$ denotes the feature dimension. Let $\boldsymbol{Y}\boldsymbol{\in}\boldsymbol{R}^{\boldsymbol{p\times n}}$ represent the feature vectors, where $\boldsymbol{Y}$ encompasses the encoder patch tokens and pre-trained decoder gene prompts, and $\boldsymbol{p}$ indicating the target feature sequence length. $\boldsymbol{Q}\boldsymbol{\in}\boldsymbol{R}^{\boldsymbol{p\times a}}$and$\boldsymbol{K}\boldsymbol{\in}\boldsymbol{R}^{\boldsymbol{m\times a}}$ represent the query and key vectors respectively, where $\boldsymbol{a}$ denotes an intermediate dimension. $\boldsymbol{V}\boldsymbol{\in}\boldsymbol{R}^{\boldsymbol{m\times b}}$represents the value vector, where $\boldsymbol{b}$ denotes the output dimension.

The cross-attention mechanism is formulated as:

$$\boldsymbol{Q}_{\boldsymbol{p\times a}}\boldsymbol{=Y}\boldsymbol{W}_{\boldsymbol{q}}\boldsymbol{,}\boldsymbol{K}_{\boldsymbol{m\times a}}\boldsymbol{=X}\boldsymbol{W}_{\boldsymbol{k}}\boldsymbol{,}\boldsymbol{V}_{\boldsymbol{m\times b}}\boldsymbol{=X}\boldsymbol{W}_{\boldsymbol{v}}$$

$$\mathbf{Attention}\left( \boldsymbol{Q,K,V} \right)\boldsymbol{=}\mathbf{softmax}\left( \frac{\boldsymbol{Q}\boldsymbol{K}^{\boldsymbol{T}}}{\sqrt{\boldsymbol{d}_{\boldsymbol{k}}}} \right)\boldsymbol{\cdot}\boldsymbol{V}$$

where $\boldsymbol{d}_{\boldsymbol{k}}$ represents the dimensionality of $\boldsymbol{K}$, and the output is a feature matrix of dimensions $\boldsymbol{p\times b}$.

**Supplementary Node 2. Self-attention Mechanism**

Let $\boldsymbol{X}\boldsymbol{\in}\boldsymbol{R}^{\boldsymbol{m\times n}}$denote the input feature vectors, where $\boldsymbol{m}$ represents the sequence length and n denotes the feature dimension. $\boldsymbol{Q}\boldsymbol{\in}\boldsymbol{R}^{\boldsymbol{m\times a}}$ and $\boldsymbol{K}\boldsymbol{\in}\boldsymbol{R}^{\boldsymbol{m\times a}}$ represent the query and key vectors respectively, where $\boldsymbol{a}$ denotes an intermediate dimension. $\boldsymbol{V}\boldsymbol{\in}\boldsymbol{R}^{\boldsymbol{m\times b}}$ represents the value vector, where $\boldsymbol{b}$ denotes the output dimension.

The self-attention mechanism is formulated as:

$$\boldsymbol{Q}_{\boldsymbol{m\times a}}\boldsymbol{=X}\boldsymbol{W}_{\boldsymbol{q}}\boldsymbol{,}\boldsymbol{K}_{\boldsymbol{m\times a}}\boldsymbol{=X}\boldsymbol{W}_{\boldsymbol{k}}\boldsymbol{,}\boldsymbol{V}_{\boldsymbol{m\times b}}\boldsymbol{=X}\boldsymbol{W}_{\boldsymbol{v}}$$

$$\mathbf{Attention}\left( \boldsymbol{Q,K,V} \right)\boldsymbol{=}\mathbf{softmax}\left( \frac{\boldsymbol{Q}\boldsymbol{K}^{\boldsymbol{T}}}{\sqrt{\boldsymbol{d}_{\boldsymbol{k}}}} \right)\boldsymbol{\cdot}\boldsymbol{V}$$

where $\boldsymbol{d}_{\boldsymbol{k}}$ represents the dimensionality of $\boldsymbol{K}$, and the output is a feature matrix of dimensions $\boldsymbol{m\times b}$, with the output dimension matching the input dimension.

This formulation preserves the structural symmetry characteristic of self-attention, where the input sequence attends to itself through learned query-key relationships.

**Supplementary Figures**

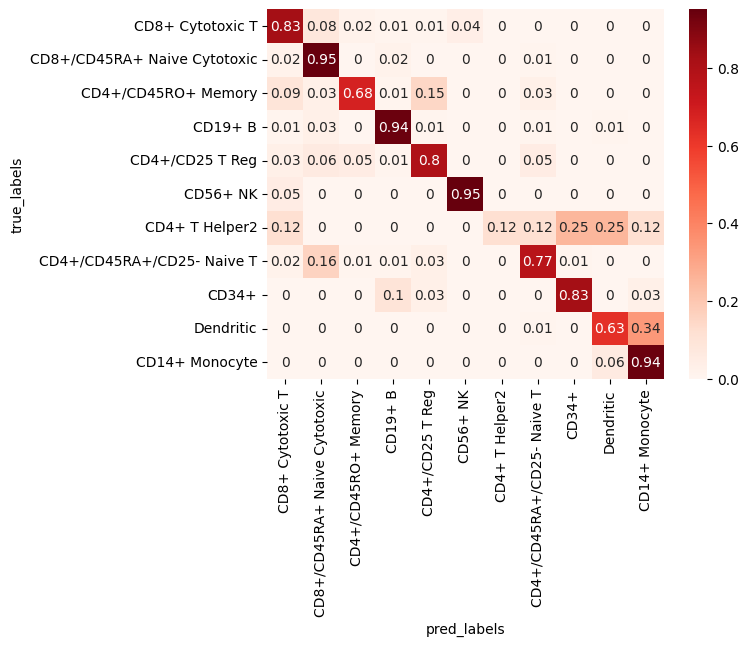

**Supplementary Figure 1. Confusion matrix analysis of CellPatch predictions on the Zheng68k dataset**. The confusion matrix displays the relationship between true cell types (*y*-axis) and predicted cell types (*x*-axis). The heatmap represents the proportion of cells from each true cell type assigned to each predicted category, with color intensity indicating the percentage of predictions. Each row sums to 100%, showing the distribution of predictions for a given true cell type.

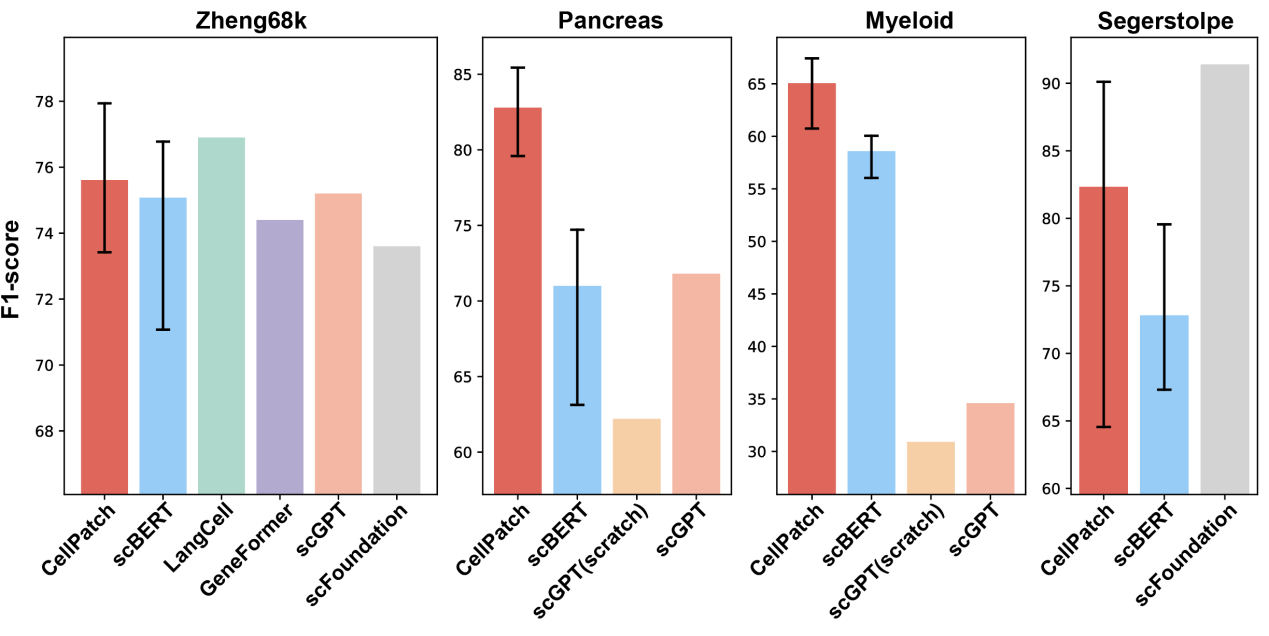

**Supplementary Figure 2. Comparative analysis of F1 scores for cell type annotation across methods and datasets.** The four subpanels demonstrate the performance of different methods on four independent datasets. Results for CellPatch and scBERT represent the mean of ten independent replicates, with error bars indicating maximum and minimum values. Results for other methods are reported values from their respective original publications.

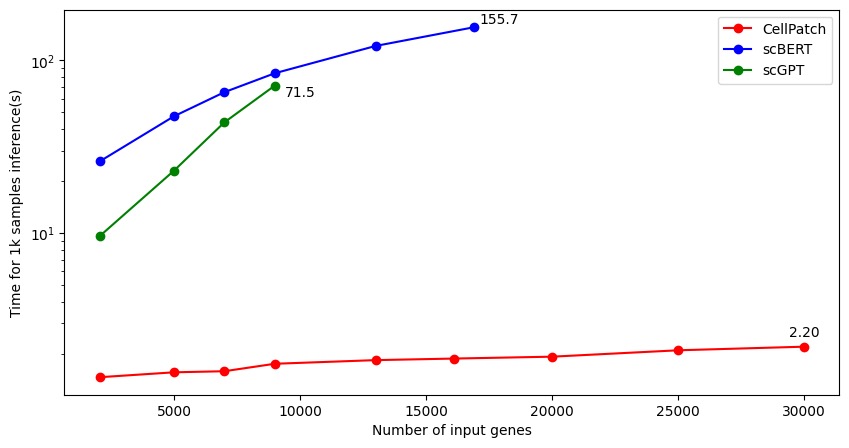

**Supplementary Figure 3. Model Speed Comparison.** The line graph illustrates the relationship between model runtime (*y*-axis) and the number of input genes (*x*-axis) across three models. The performance of the models was evaluated on a simulated cell type annotation task, with 1000 cells simulated as input for each model. For each model, the runtime was measured across various number of input genes: 2048, 5k, 7k, 9k, 13k, 16906, 20k, 25k, and 30k (corresponding to the y-axis). Here, 2048 is the default number of input genes for scGPT, and 16906 is the default number of input genes for scBERT. All calculations were performed on an A100 80G GPU under the same conditions. The batch sizes for CellPatch, scBERT, and scGPT were 128, 4, and 3, respectively. Notably, scBERT ran out of GPU memory at 20k or more genes as input, while scGPT ran out of GPU memory at 13k or more. In contrast, CellPatch was able to handle up to 30k genes as input without exceeding GPU memory limits. Given that a feature number of 30k is sufficient for vast majority of single-cell RNA sequencing data requirements, including those for multimodal inputs, further testing with larger number of features was not conducted. CellPatch exhibits significantly lower time consumption than scBERT and scGPT, while achieving comparable or even superior performance.

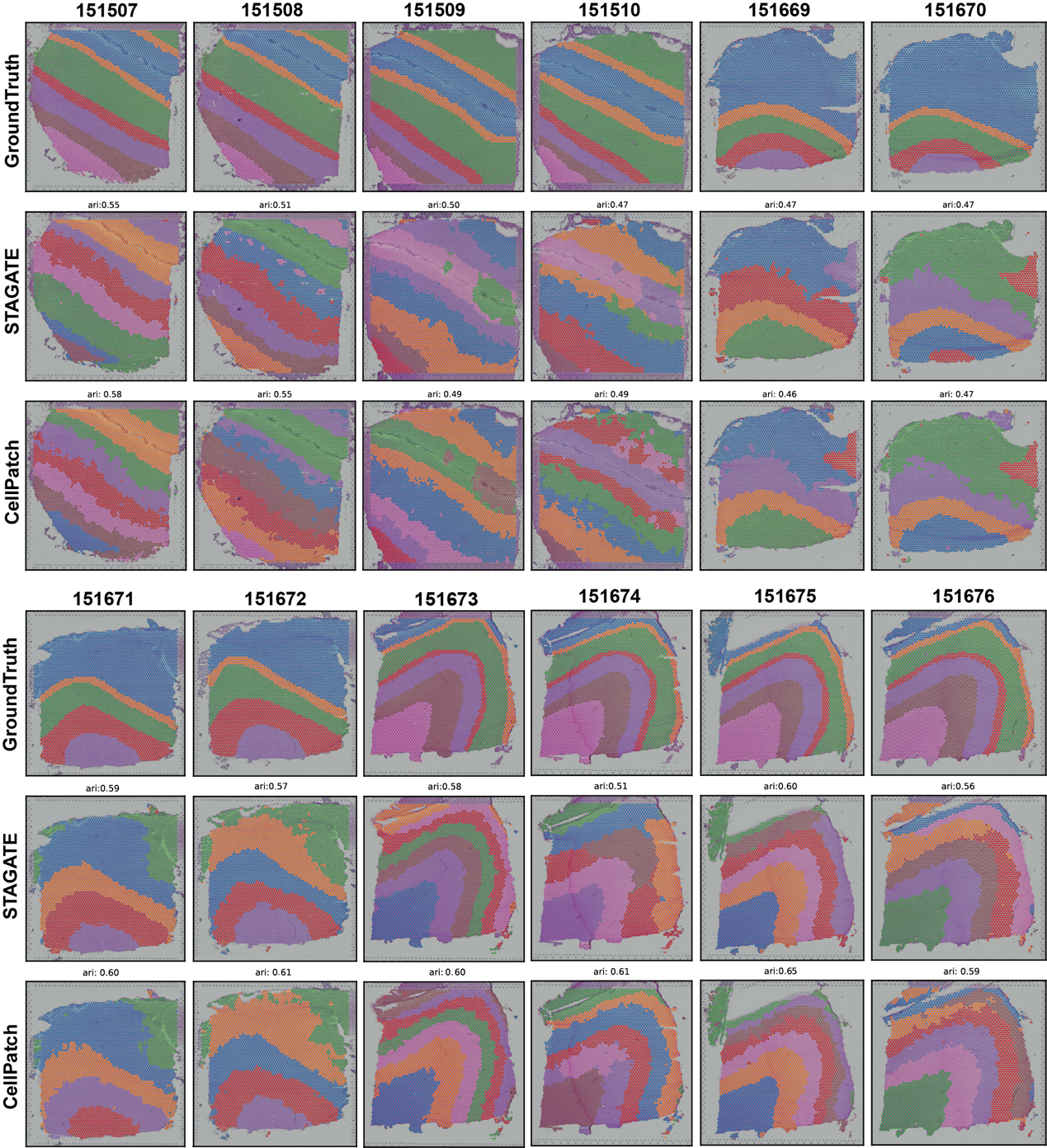

**Supplementary Figure 4. Spatial transcriptomic clustering results on the DLPFC dataset.** The figure presents spatial clustering results for 12 samples from the DLPFC dataset. Each sample is represented by three spatial clustering maps arranged vertically: ground truth annotation (top), original STAGATE clustering results (middle), and STAGATE results enhanced by CellPatch (bottom, labeled as 'CellPatch').

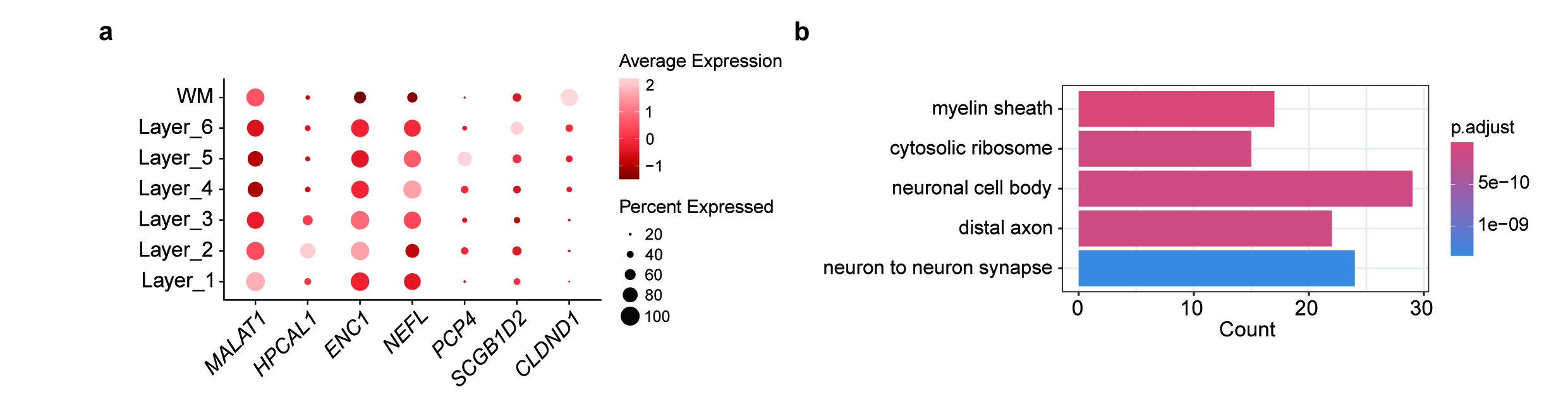

**Supplementary Figure 5. Downstream analysis on sample 151675 of the DLPFC dataset.** **a,** Scatter plot displaying the mean expression of layer-specific markers including *CLDND1* (white matter), *SCGB1D2* (layer 6), *PCP4* (layer 5), *NEFL* (layer 4), *ENCL* (layer 3), *HPCAL1* (layer 2), and *MALAT1* (layer 1). Clusters in *y*-axis correspond to the spatial domains inferred by CellPatch. **b,** Gene Ontology (GO) and Kyoto Encyclopedia of Genes and Genomes (KEGG) biological pathway analysis for the layer-specific marker genes detected by CellPatch on the sample 151675. Enrichment is calculated as -log10 adjusted p-value of the layer-specific marker genes.

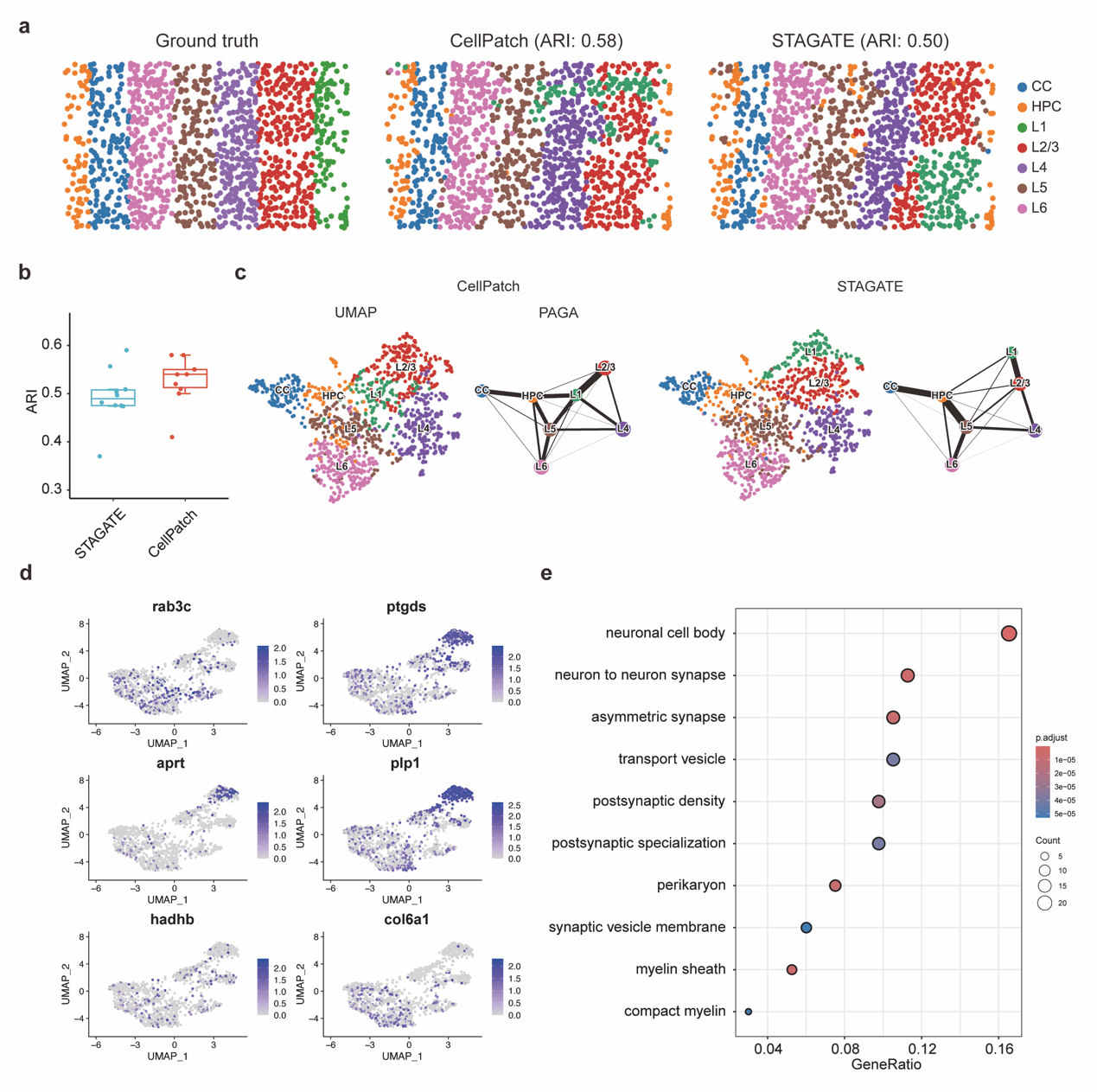

**Supplementary Figure 6. Performance of CellPatch on mouse medial prefrontal cortex data by STARmap.** **a,** Manual annotation and spatial domain detection results of CellPatch and STAGATE on mouse medial prefrontal cortex data. **b,** boxplot illustrating clustering accuracies across 10 repeats. **c,** UMAP visualizations and PAGA graphs of STARmap data generated from CellPatch and STAGATE embeddings, respectively. **d,** UMAP visualizations of layer-specific marker genes expression. **e,** Gene Ontology (GO) and Kyoto Encyclopedia of Genes and Genomes (KEGG) biological pathway analysis for the layer-specific marker genes detected by CellPatch. Enrichment is calculated as -log10 adjusted p-value of the layer-specific marker genes.

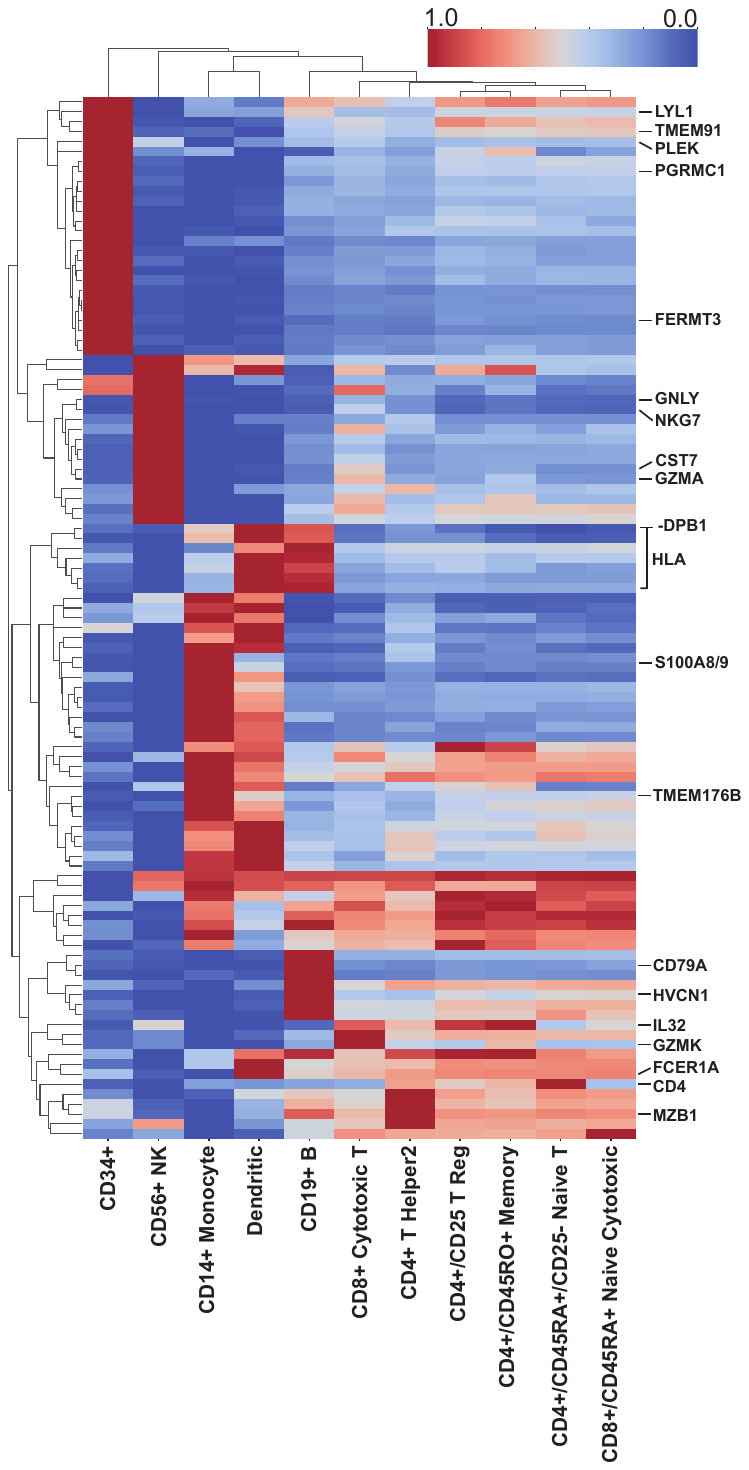

**Supplementary Figure 7. Cell type-specific attention score heatmap of differentially expressed and marker genes in the Zheng68k dataset.** The heatmap displays first-layer attention scores across cell types (*x*-axis) for a curated set of genes (*y*-axis), comprising 30 differentially expressed genes and established markers per cell type in the Zheng68k dataset. Color intensity represents the magnitude of attention scores, revealing cell type-specific gene recognition patterns.

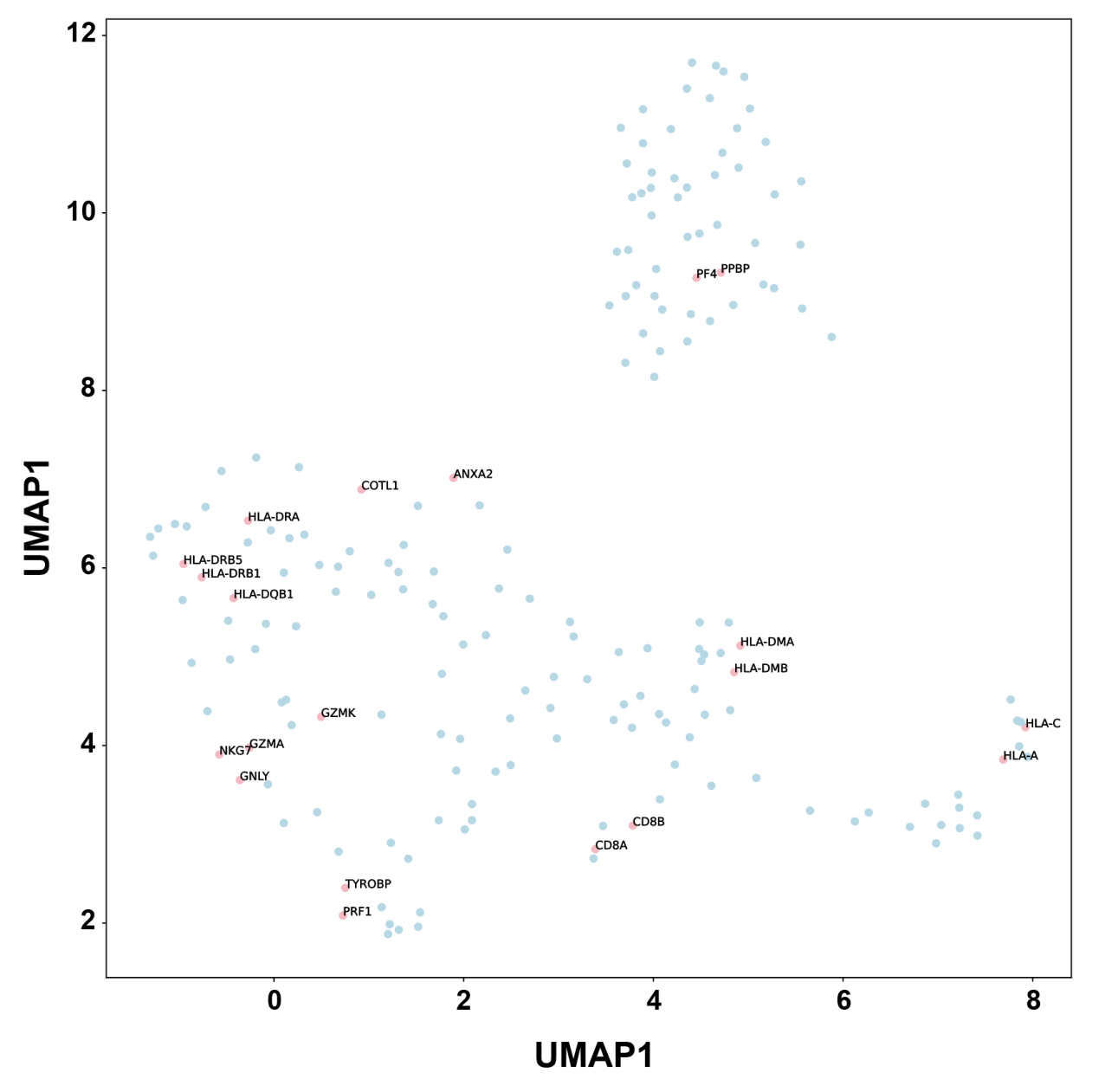
**Supplementary Figure 8. UMAP visualization of gene embeddings learned by CellPatch from the Zheng68k dataset.** The plot visualizes the UMAP projection of gene embeddings extracted from CellPatch, featuring 30 differentially expressed genes and established markers for each cell type in the Zheng68k dataset. Each point represents the two-dimensional UMAP projection of a gene's embedding vector. The spatial clustering of functionally related genes demonstrates CellPatch's capability to capture meaningful gene-gene relationships through its embedding space.

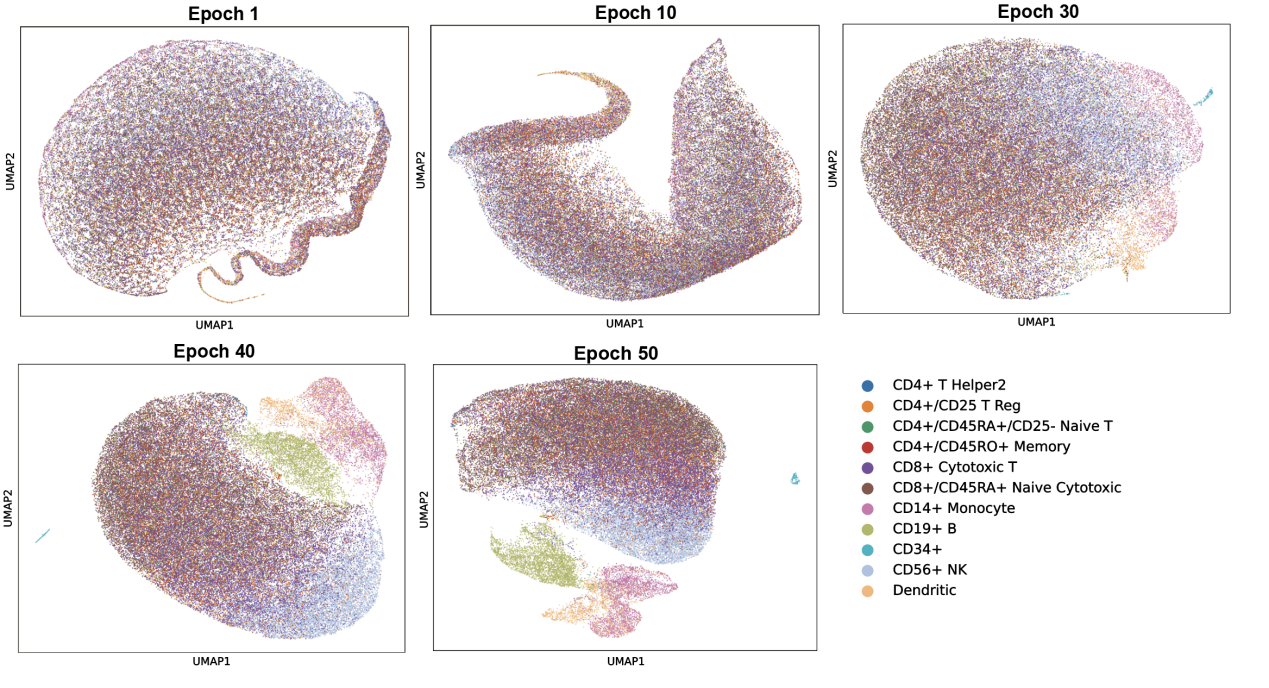

**Supplementary Figure 9. UMAP visualization of Zheng68k dataset features across pre-training epochs.** The panel displays UMAP projections of cell features extracted from CellPatch at epochs 1, 10, 30, 40, and 50 of pre-training. Cell types are distinguished by different colors. The progressive separation of cell type clusters demonstrates CellPatch's increasing discriminative power throughout the pre-training process.
